# Joint evolution of mycorrhizal type, pollination, and seed dispersal mode in trees

**DOI:** 10.1101/2020.10.04.325282

**Authors:** Akira Yamawo, Misuzu Ohno

**Affiliations:** Department of Biological Sciences, Faculty of Agriculture and Life Science, Hirosaki University, 3 Bunkyo-cho, Hirosaki, Aomori 036-8561, Japan

## Abstract

Although tree diversity is fundamental to terrestrial ecosystems, the processes that generate it remain uncertain. Mycorrhizal type, pollination mode, and seed dispersal mode may be key drivers of tree diversity. We predicted that mycorrhizal symbiosis drove evolution of pollination and seed dispersal modes because arbuscular mycorrhizal (AM) associations would favour long-range seed or pollen dispersal owing to their negative plant–soil-feedback effects on conspecific individuals, and ectomycorrhizal (EcM) associations would favour short-range dispersal owing to their positive effects. We analysed evolutionary relationships among mycorrhizal type, seed dispersal mode, and pollination mode in 704 tree species and conducted a meta-analysis of the dispersal distances of the various seed dispersal and pollination modes. We found evidence of joint evolution of all three features. Most AM-associated trees had endozoochorous seed dispersal and biotic pollination and these dispersal modes had long dispersal distances, whereas most EcM-associated trees had anemochorous seed dispersal and wind pollination and these dispersal modes had relatively shorter dispersal distances. Overall, evolution of mycorrhizal type, seed dispersal mode, and pollination mode were linked, strongly suggesting that mycorrhizal symbiosis drives the evolution of seed and pollination modes and contributes to tree diversification.

## Introduction

Tree diversity provides the energetic, structural, and ecological basis of many terrestrial ecosystems. However, the processes that underlie its formation remain uncertain. Mycorrhizal symbiosis, pollination, and seed dispersal are major factors that characterise tree diversity. Mycorrhizal symbiosis is the oldest known biological interaction in plants: plants have lived closely with mycorrhizal fungi for over 400 million years (Tedersoo et al., 2020). Trees have evolved also several types of mycorrhizal symbiosis such as arbuscular (AM), ectomycorrhizal (EcM) and ericoid. The mycorrhizal symbiosis is recognised as an important factor in growth and survival of trees and maintaining the diversity of tree species in forest ecosystems (Tedersoo et al., 2020; Teste et al., 2017). Mycorrhizal fungi provide their host plants with nutrients from the soil or aid in defence against insects and parasitic fungi and enhance the survival rate, growth, flower production, and fitness of their host plants (Tedersoo et al., 2020). Pollination modes include biotic and wind for over 400 million years (Wang et al., 2019). Biotic pollination depends on other organisms such as insects, bats, and birds, whereas wind pollination depends primarily on wind. Seed dispersal modes include biotic (endozoochory), wind (anemochorous), and unspecialised (Heleno & Vargas, 2015). Endozoochory depends on a wide taxonomic variety of organisms, such as mammals, birds, and insects. Anemochorous and unspecialised seed dispersal depend on wind or gravity. Thus, trees have evolved various mycorrhizal symbiosis, pollination, and seed dispersal modes, however, their evolutionary interactions have not been elucidated.

Mycorrhizal fungi contribute to the maintenance of tree community diversity via plant–soil feedbacks (PSFs), which can be either positive or negative and which differ between AM and ectomycorrhizal EcM fungi (Bennett et al., 2017; Tedersoo et al., 2020; Teste et al., 2017). The effects of these different PSFs promote variation in population density in tree species (Bennett et al., 2017), and differences in PSFs and population density may drive the evolution of pollination or seed dispersal. Biotic pollen and seed dispersal may permit greater dispersal distances than wind or unspecialised dispersal (Hesse & Pannell, 2011; Mustajärvi et al., 2001; Pellmyr, 1992). Thus, negative PSFs may favour biotic seed dispersal as long-range seed dispersal in AM-associated trees, and positive PSFs may favour wind or unspecialised seed dispersal as short-range seed dispersal in EcM-associated trees. Because negative and positive PSFs may favour low and high population densities (Bennett et al., 2017), respectively, they are expected to favour biotic pollination, enabling dispersal over long distances (Pellmyr, 1992), and wind pollination, effective over short distances (Hesse & Pannell, 2011; Mustajärvi et al., 2001), respectively. However, mycorrhizal type, seed dispersal mode, and pollination mode have traditionally been studied in isolation from each other, and their combined effects remain a major gap in our understanding of tree evolution. Given the ubiquity of interactions with mycorrhizal fungi, pollinators, and seed dispersers among tree species, it is surprising how little is known of the evolutionary relationships among these interactions.

Although there are several studies of the effects of mycorrhizal fungi on pollination, most studies are limited to experimental approaches focused on ecological effects in a few species (Tedersoo et al., 2020; Wolfe et al., 2005; but see Correia et al., 2018). A phylogenetic approach that includes multiple species can close this gap by directly demonstrating the evolutionary patterns among these interactions (Correia et al., 2018). We conducted a meta-analysis of seed and pollen dispersal distance in each mode, and investigated the evolutionary relationships among mycorrhizal type (AM, EcM, and both AM + EcM), seed dispersal (by animals: endozoochorous; by wind: anemochorous; and unspecialised) and pollination mode (biotic and wind) in 704 tree species from different regions throughout the world, including Asia, America, Africa, Europe and Oceania, in three climate areas (tropical, subtropical and temperate), by phylogenetic analysis.

## Materials and Methods

### Phylogenetic relations among mycorrhizal type, pollination, and seed dispersal mode

We collected information on mycorrhizal type, seed dispersal mode, and pollination mode from multiple databases. Species for which all three data were not available were excluded.

#### Mycorrhizal types

Six main sources were used to compile information on tree mycorrhizal type (AM, EcM or AM + EcM): (1) Mycoflor, a database that lists the mycorrhizal status of tree species (Hempel et al., 2013); (2) an updated checklist of mycorrhizae of land plants (Wang & Qiu, 2006); (3) a checklist of mycorrhizae of the flora of Britain (Harley & Harley, 1987, 1990); (4) a database documenting mycorrhizal associations in 3000 vascular plant species from the former Soviet Union (Akhmetzhanova et al., 2012); and (5) a dataset from the Global Biodiversity Information Facility (https://www.gbif.org) (Soudzilovskaia et al., 2020). Other sources of information on mycorrhizal type are included in the Supporting Information. To minimise errors, the data were individually checked in multiple databases and their cited references.

#### Primary seed dispersal and pollination modes

Data on seed features were collected from the list compiled by Heleno and Vargas (2015), the Plants of the World Online (POWO; http://www.plantsoftheworldonline.org/), and the Global Biodiversity Information Facility (Edwards et al., 2000) (Supporting Information). The list compiled by Heleno and Vargas (2015) includes the LDD characteristics of 10,792 species from 142 families in the *Flora Europaea* (Tutin et al., 1980). POWO is a Web portal that makes available scientific data collected at the Royal Botanical Gardens, Kew, including the names of over a million plants from around the globe, plus images and detailed descriptions of many of those. Seeds with characteristics that promote wind dispersal, such as wings or a pappus, were classified as anemochorous; seeds with tissues that attract animals, such as fleshy or nutritious tissues, were classified as endozoochorous (Edwards et al., 2000; Vargas et al., 2012). Other species were classified as unspecialised.

The pollination mode was classified as either biotic or wind (Wang et al., 2019), and our classification scheme did not include other categories, such as self-pollination. Flowers with typically developed petals were classified as having a biotic pollination mode. Flowers without petals were classified as wind pollinated. Information for the classification of pollination modes was collected from the above databases (Supporting Information).

#### Phylogenetic tree

Plant List 1.1 was used to standardise species names, and membership in higher taxonomic groups was standardised against Angiosperm Phylogeny Group IV (Angiosperm Phylogeny Group, 2016). Our database contains information on the mycorrhizal types, seed dispersal, and pollination modes of 704 plant species from 343 genera in 100 families. The most represented families in our study were the Leguminosae (14.5%), Rosaceae (8.5%), Pinaceae (8.0%), Fagaceae (5.0%) and Betulaceae (3.3%).

We used PhytoPhylo tree software (Qian & Jin, 2016) to construct a phylogenetic tree based on the phylogeny generated by Zanne et al. (2014) and updated by Qian and Jin (2016). Then we reconstructed this tree to incorporate the phylogenetic correction based on the PSR index by using the ‘S.PhyloMaker’ function implemented in R. The final phylogenetic tree had 704 tip labels and 677 internal nodes (Fig. 1).

**Fig. 1.**
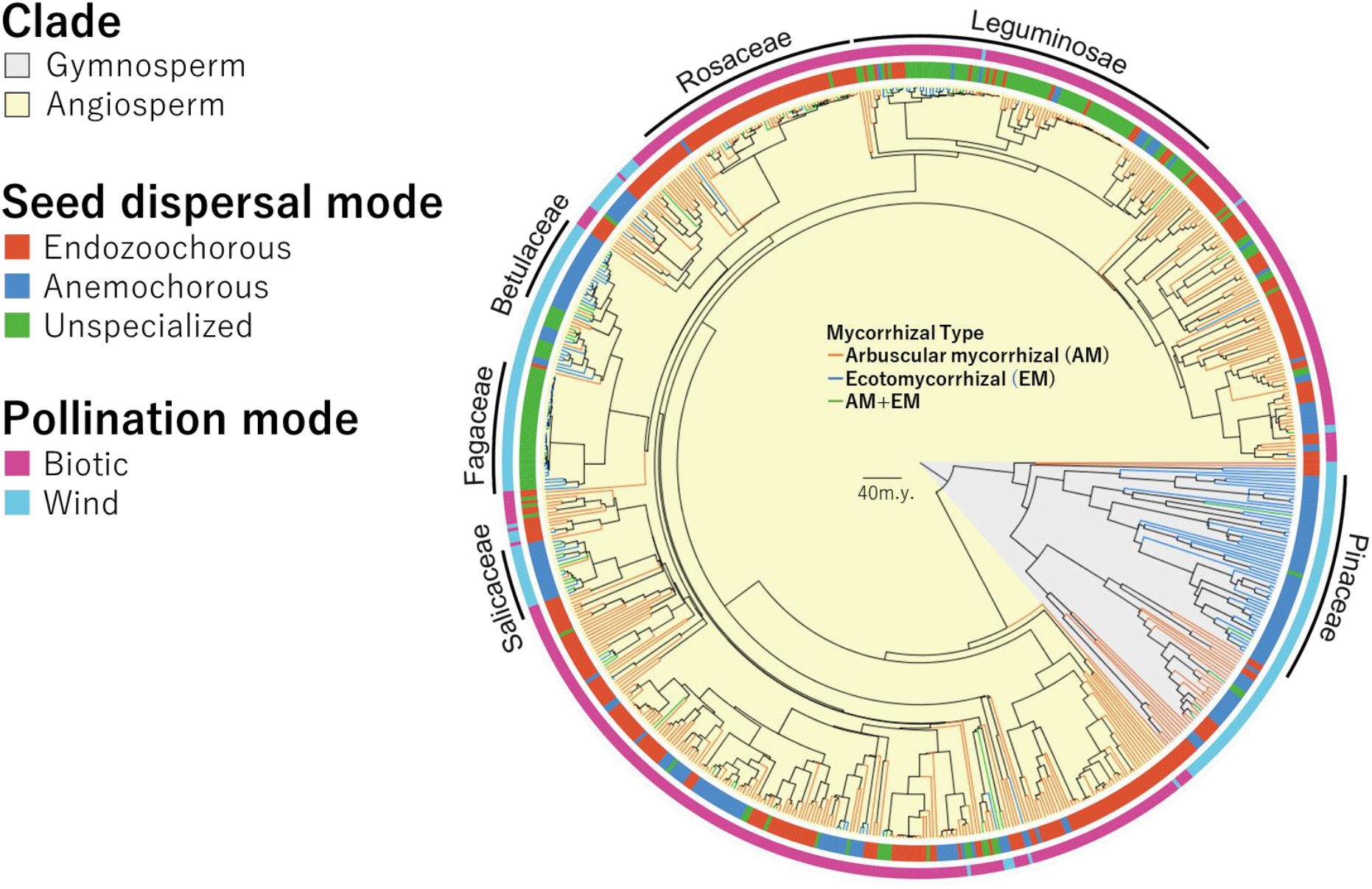
Time-calibrated phylogeny of the 704 plant species used in this study. Branch colour denotes mycorrhizal type (orange, AM; blue, EcM; green, AM + EcM). Colours of inner and outer labels ringing the tree indicate seed dispersal mode (red, endozoochorous; blue, anemochorous; green, unspecialised) and pollination mode (pink, biotic; blue, wind). The scale bar represents 40 million years.

### Meta-analysis of seed and pollen dispersal distances

We searched the Web of Science (http://apps.webofknowledge.com) for articles in English and Google Scholar (https://scholar.google.com/) for papers in English and Japanese that included ‘tree’, ‘woody plant’, ‘seed dispersal distance’, or ‘pollination distance’ in the title, abstract, or keywords. All peer-reviewed papers published between 1970 and 2020 that investigated maximum and/or mean value of seed dispersal distance or pollination distance were selected and examined. We inspected their references and added studies not found in the initial search. Dispersal mode were divided based on information of each article. Data were compiled following rules. First, we used only studies based on naturally occurring species in the field; we excluded studies that used artificial seeds or artificial modified diaspores.

Second, to ensure that the dispersal distances were accurate, we selected only studies that included direct measurements of seed or pollen dispersal distance or direct estimates of the distance travelled by dispersers using genetic markers. In other words, we did not include studies that estimated dispersal distance by using models, because dispersal distances estimated by using models are higher than those estimated based on experiments (Chen et al., 2019). In cases where data were available only as figures, these were digitised, and the maximum and mean values was determined in Getdata Graph Digitizer software (http://www.getdata-graph-digitizer.com/). We combined seed dispersal distance data of tree plant species from available global syntheses (Chen et al., 2019) to create our database. We remove data of ballistic seed dispersal mode, shrub and herb plants. Abiotic seed dispersal was divided into anemochorous and unspecialised based on seed morphology (Heleno &Vargas, 2015). When data for the same dispersal mode in a given species were available in multiple studies, we used the mean value for analysis. The search, completed on 23 January 2021, produced 186 peer-reviewed scientific studies (maximum seed dispersal of 187 species from 131 genera in 53 families, mean seed dispersal of 145 species from 108 genera in 45 families, maximum pollination dispersal distance of 82 species from 68 genera in 39 families and mean pollination dispersal distance of 36 species from 33 genera in 25 families); these studies were then used for the meta-analysis (Supporting Information). Secondary seed dispersal distances were also collected from the dataset published in Chen et al. (2019) and web searches using the rules mentioned above (96 peer-reviewed scientific studies) (maximum seed dispersal distances of 87 species from 51 genera in 31 families and mean seed dispersal distances of 94 species from 51 genera in 30 families), and compared with primary seed dispersal distances. For all analyses of seed and pollen dispersal distances, we used both maximum and mean values.

#### Statistical analyses

##### Phylogenetic relations among mycorrhizal type, pollination mode, and seed dispersal mode

Log–linear analysis with maximum likelihood chi-squared and post hoc Freeman–Tukey deviation tests, with Bonferroni’s correction, were used to test for univariate relationships between mycorrhizal type, seed dispersal, and pollination modes (Legendre & Legendre 1998). All analyses were performed in R v. 4.1.0 software (R Development Core Team, 2021).

In all analyses, a phylogenetic species richness (PSR) index was used to correct for phylogenetic relatedness (Helmus et al., 2007). To calculate PSR, the number of species in a sample is multiplied by the phylogenetic species variability, a term that describes how the variance of a trait is decreased by the phylogenetic relatedness of the species being studied (Helmus et al., 2007). PSR thus represents the species richness of a sample after species relatedness has been discounted, making it more appropriate than uncorrected species richness for use in the contingency table. PSR values decrease as species relatedness increases, approaching zero (Helmus et al., 2007).

First, mycorrhizal type (AM, AM + EcM, and EcM) was tested against the proportion of seed dispersal mode (anemochorous, endozoochorous, and unspecialised) or pollination mode (biotic and wind). Seed dispersal mode was also tested against the proportion of pollination mode. For each pair of mycorrhizal type and/or dispersal mode, PSR was calculated in the R package ‘picante’ (Kembel et al., 2010); these values were used in the contingency tables as described in Pyšek et al. (2011) and Hempel et al. (2013).

Contingency tables were used to check for differences between categories by use of log– linear models with maximum likelihood chi-square analysis. Post-hoc Freeman–Tukey deviation tests (Sokal & Rohlf, 1995) with Bonferroni’s correction were used for two-way or multiway contingency tables. In the Freeman–Tukey test, the observed and expected counts were used to calculate an approximately normalised estimate of how far each observed frequency deviates from the null hypothesis (Freeman–Tukey deviate: √ (O) + √(O + 1) – √(4E + 1)). We also used the approach described by Sokal and Rohlf (1995) to calculate a critical value for the Freeman–Tukey deviates (√(*v* X _[1,α]_ ^2^ / (no. cells)).

##### Meta-analysis of seed and pollen dispersal distances

All data were transformed to log10, to secure normality distribution. Maximum and mean dispersal distance of seed or pollen among the dispersal modes were compared used a phylogenetic Generalized Least Squares analysis (pGLS) (Paradis, 2011) by the gls function in the nlme package in R version 4.1.0 (R Development Core Team, 2021) to account for evolutionary history of the species (Felsenstein, 1985). The plant phylogenetic tree based PhytoPhylo tree software (Qian & Jin, 2016) were used for pGLS. We used Pagel’s λ correlation structure (Revell, 2010), to incorporate the information of phylogeny into the models. The corPagel function in the ape package (Paradis, 2019) with the amount of phylogenetic signal in the data estimated using restricted maximum likelihood (REML) approach was used for constructing the models (Freckleton et al., 2002). Dispersal distances of seeds or pollen were include as response variable, and seed dispersal modes (endozoochorous, anemochorous, or unspecialised) or pollen dispersal modes (biotic or wind) were included as explanatory variables. To reveal the effects of secondary seed dispersal, secondary seed dispersal distance also compared with primary seed dispersal distance in each dispersal mode. *P*-values were corrected by the holm method.

## Results and Discussion

All trees (704 plant species from 343 genera in 100 families) in our database were assigned to one of three mycorrhizal types: AM (480 species from 295 genera in 94 families), AM + EcM (62 species from 35 genera in 20 families), or EcM (162 species from 53 genera in 18 families). Association with AM fungi was widespread across the phylogenetic tree, being present in ∼90% of all families and ∼97% of genera (Fig. 1). Both angiosperm and gymnosperm clades were present. Tree species of all three mycorrhizal types were present in both clades. Similarly, seed dispersal modes (endozoochorous, 333 species from 187 genera in 81 families; anemochorous, 213 species from 78 genera in 30 families; unspecialised, 158 species from 88 genera in 24 families; Fig. 1) and pollination modes (biotic, 485 species from 279 genera in 83 families; and wind, 219 species from 67 genera in 26 families) were scattered across the phylogenetic tree.

The phylogenetically corrected log–linear analysis revealed that mycorrhizal types and seed dispersal modes were significantly linked (Table 1). Of the AM-associated trees, a greater proportion produced endozoochorous seeds and lower proportion produced anemochorous seeds compared with number of species than expected by the log–linear model (Fig. 2A; Table S1). This result was not found in the analysis by Correia et al. (2018) that included both tree and herb species, which indicates that it is specific to trees and their life-history traits. In contrast, more EcM-associated trees produced anemochorous seeds and lower proportion produced endozoochorous seeds compared with number of species than expected by the log–linear model (Fig. 2A; Table S1). Our meta-analysis revealed that both the maximum and mean dispersal distances of endozoochorous seeds are longer than those of anemochorous seeds and unspecialised seeds (Fig. 3A and B, Table S2). Although several studies reported secondary dispersal (e.g., by mice or ants) in anemochorous and unspecialised seeds, the secondary seed dispersal distances were shorter than the endozoochorous and anemochorous dispersal distances, and did not differ from the dispersal distances of unspecialised seeds (Table S3). Thus, endozoochorous dispersal is more effective for long-distance seed dispersal than other dispersal modes. Several studies have reported that within-species PSFs differ between AM- and EcM-type trees (Tedersoo et al., 2020). For EcM-associated trees, PSFs are positive within species: growth and survival rate are increased in soil in which conspecifics grow (Bennett et al., 2017). Conversely, for AM-type trees, PSFs are negative within species: growth and survival rate are reduced in soil in which conspecifics grow (Bennett et al., 2017). These contrasting effects should promote the evolution of different seed dispersal modes in AM- and EcM-type trees. EcM-associated trees may have evolved middle-range seed dispersal by wind (Figs. 2A, 3A and B) because it allows seedlings to receive positive PSFs from conspecifics but avoid competition with parental trees. In contrast, AM-associated trees may have evolved long-range seed dispersal by animals (Figs. 2A, 3A and B), because it allows seedlings to avoid negative PSFs from conspecifics.

**Table 1.** Results of phylogenetically corrected log-linear analyses examining relationships between mycorrhizal type, seed dispersal, and pollination mode in 704 tree species.

|  | Seed dispersal mode | Pollination mode |
| --- | --- | --- |
| Mycorrhizal type | $N = 704, df = 4,$<br>$\chi^2 = 94.61, P < 0.001$ | $N = 704, df = 2,$<br>$\chi^2 = 126.18, P < 0.001$ |
| Seed dispersal mode | | $N = 704, df = 2,$<br>$\chi^2 = 108.13, P < 0.001$ |

**Fig. 2.**
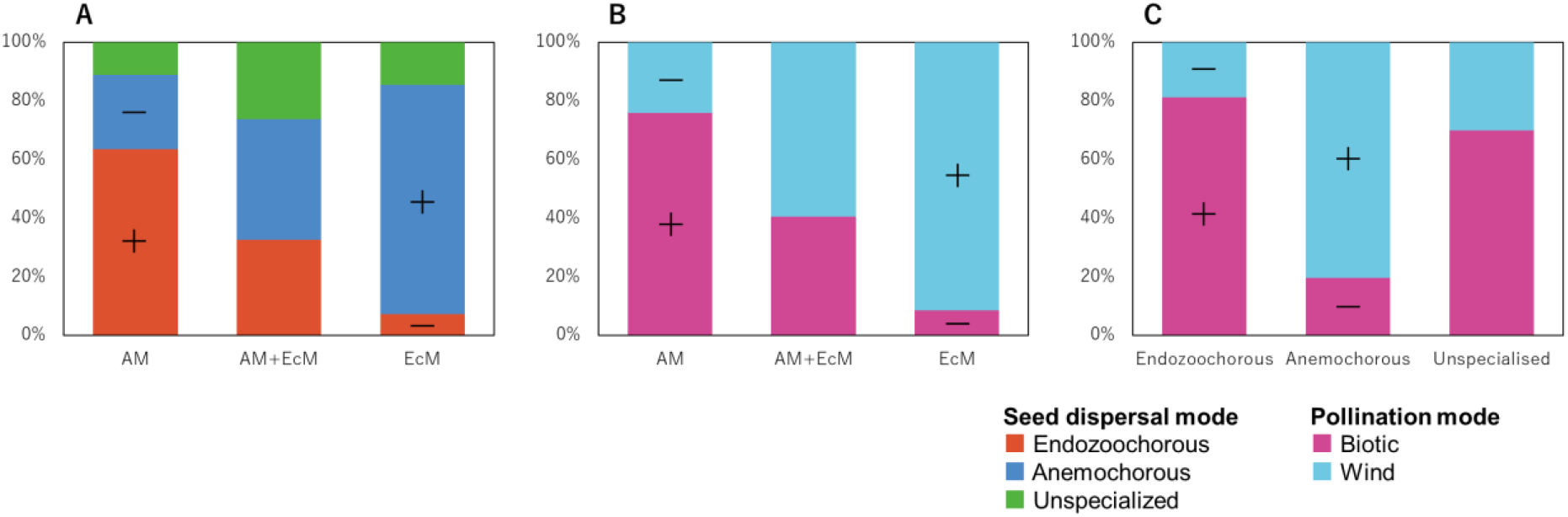
(A) Relative proportions (% of total) of specialised structures for seed dispersal mode in trees with different mycorrhizal types: AM, AM + EcM, EcM. (B) Relative proportions (% of total) of pollination modes in trees with different mycorrhizal types. (C) Relative proportions (% of total) of pollination modes in trees with different seed dispersal modes: biotic and wind. Plus (+) signs indicate significantly higher percentages, and minus (−) signs indicate significantly lower percentages than expected by the log–linear model (Freeman– Tukey deviation test, *P* < 0.001).

**Fig. 3.**
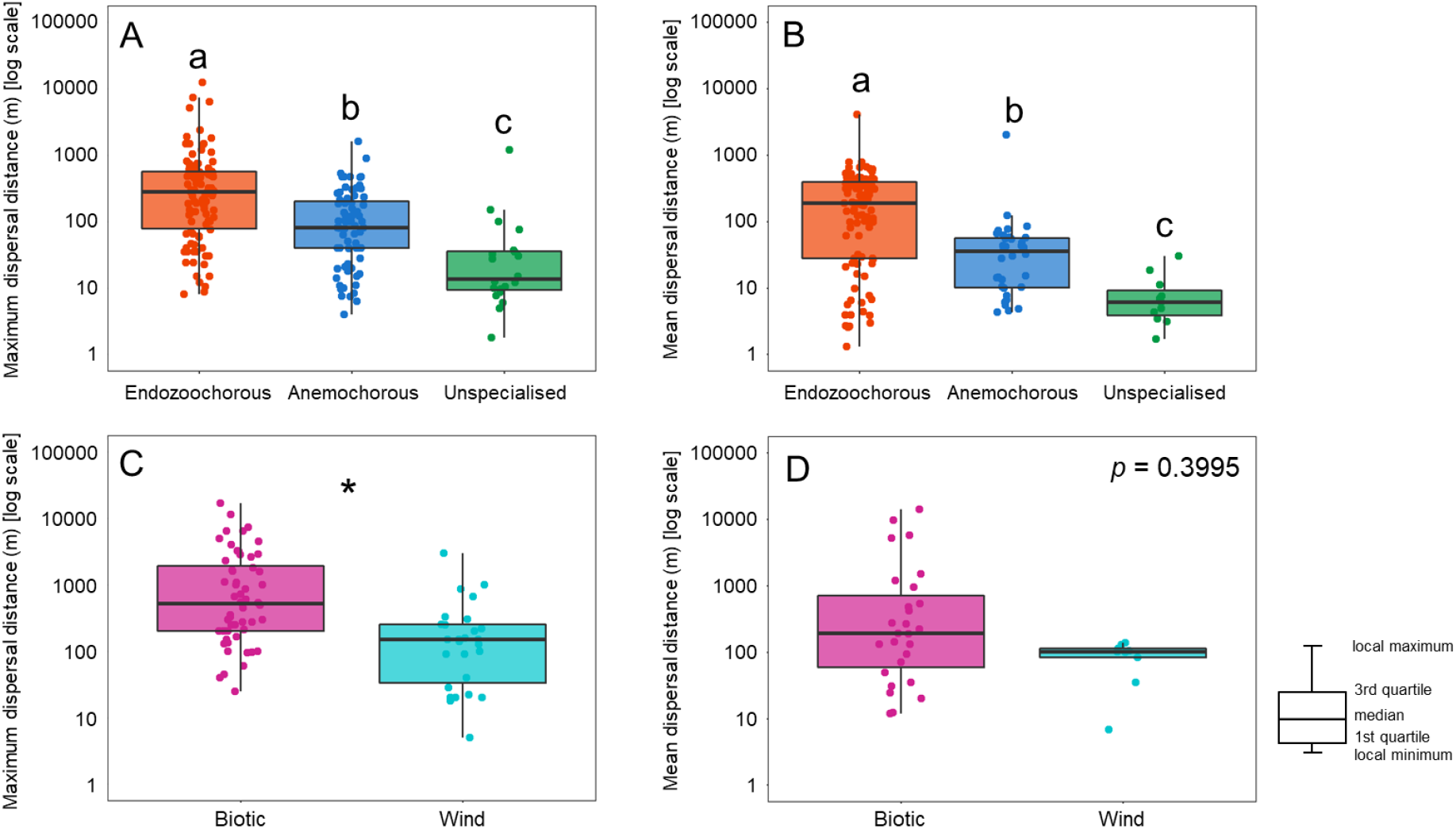
(A) Maximum and (B) mean seed dispersal distance of each seed dispersal mode. (C) Maximum and (D) mean pollen dispersal distance of each pollination mode. Different letters and asterisks indicate that values differ significantly at \**P* < 0.05 (phylogenetic Generalized Least Squares analysis [pGLS] [Paradis 2011]).

The differences in seed dispersal modes can also be explained with respect to the different ecological characteristics of different mycorrhizal types. Mycorrhizal interactions are largely generalist, but EcM fungi are more host-specific than AM fungi (van der Heijden et al., 2015). Long-distance seed dispersal may favour symbiosis with generalist AM fungi over that with host-specific EcM fungi through higher encounter rates with generalist AM fungi around them after dispersal. These ecological conditions also favour the evolution of different seed dispersal modes between AM- and EcM-type trees. These processes may have facilitated the joint evolution of mycorrhizal types and seed dispersal modes by interacting, not exclusively, with each other.

Pollination mode was also associated with mycorrhizal type (Fig. 2B; Tables 1, S4) and seed dispersal mode (Fig. 2C; Table 1, S5). Approximately 76% of AM-type plants are biotically pollinated, whereas more than 91% of EcM-type trees are wind-pollinated. Approximately 81% of endozoochorous seed type plants are biotically pollinated, whereas more than 80% of anemochorous seed type plants had wind-pollinated. Thus, the evolution of pollination modes may have been driven by mycorrhizal types and/or seed dispersal modes. The negative PSFs promoted by AM fungal associations and the positive PSFs promoted by EcM fungal associations with their linkage to different seed dispersal modes induce low (AM) and high (EcM) tree population densities (Bennett et al., 2017; Vittoz & Engler, 2007). Wind pollination is more effective when plants have a high population density, and the efficacy decreases as population density decreases because the pollen concentration in the air rapidly decreases with the distance from the pollen-producing plants (Culley et al., 2002). Moreover, the maximum pollen dispersal distance for wind pollination was significantly shorter than that for biotic pollination (λ= 0.718, *F* = 5.36, *P* = 0.02; Fig. 3C), but the mean dispersal distance did not differ significantly (λ= 0.764, *F* = 0.73, *P* = 0.40; Fig. 3D). Therefore, many tree species that are wind-pollinated, such as those in the Pinaceae and Fagaceae, often have high population density and are the dominant trees in the forest (Culley et al., 2002). The significantly longer maximum pollen dispersal distance for biotic pollination (Fig. 3C, Table S2) is achieved through the foraging behaviour of pollinators (Nason et al., 1998; Kremer et al., 2012). Biotic pollination thus supports effective pollination of AM-type trees with low population densities. Previous studies considered that pollination limitation and wind environment (windy or non-windy) are important factors in the evolution of wind and biotic pollination modes (Culley et al., 2002). Future studies in pollination biology should investigate the relationships between plant population density and mycorrhizal symbiosis to understand the evolution of pollination modes in trees.

By contrast, gymnosperms, which have AM-type fungal associations, did not develop biotic pollination (Fig. 1). This may be due to a phylogenetic constraint, because gymnosperms originated before the evolution of biotic pollination (Kenrick & Crane, 1997). Overall, the evolution of pollination mode appears to be linked to the evolution of mycorrhizal type via the population density of each plant species.

Our findings provide novel perspectives for understanding the evolution of mycorrhizal types and pollination and seed dispersal modes. Nevertheless, the categorisation of seed dispersal and pollination modes was based on seed and flower traits, which might not completely reflect their actual functions. For example, certain *Salix* tree species, which do not develop petals, depend on insects for pollination (Karrenberg et al., 2002). Despite these limitations, our results will facilitate our understanding of the relationships among different traits and their evolution, because in many cases, traits reflect pollination and seed dispersal functions (Correia et al., 2018; Culley et al., 2002). Future research on the detailed ecology of each tree species could improve our understanding of the processes that generate tree diversity. For example, although *Salix* tree species belong to a large EcM-associated clade, some *Salix* species are associated with AM mycorrhizae (Fig. 1). The dependence of some Salix tree species on insects for pollination may reflect changes in mycorrhizal types, and it may represent an intermediate stage in the evolution of a new pollination mode in association with changes in mycorrhizal type. Thus, tree species with mycorrhizal associations that do not fit the patterns shown here may present excellent opportunities for future research.

Plants have lived in close association with AM fungi for over 400 million years. Fossils of the earliest land plants provide evidence that fungi morphologically similar to the extant Glomales (AM mycorrhizal) lived within their cells (Heckman et al., 2001; Remy et al., 1994). Biotic pollination and endozoochory have sometimes been considered as key characteristics that led to the evolutionary success of the angiosperms beginning in the Lower Cretaceous 130 million years ago (Crepet, 1984; Pellmyr, 1992; Regal, 1977). We do not know whether these ancient AM-type trees had negative PSFs on conspecific individuals, but if they had, PSFs may have driven the evolution of pollination and seed dispersal mutualism in tree species. This raises a novel question: at what point in evolutionary time did negative PSFs appear and how did they contribute to plant diversification, together with the evolution of seed dispersal and pollination? This question may be clarified by elucidating the detailed mechanisms of negative PSFs and understanding the evolutionary history of the microbes involved.

Our findings also explain the latitudinal gradient of seed dispersal and pollination modes reported by previous studies of AM- and EcM-type trees. The greater prevalence of endozoochory in tropical regions (Wandrag et al., 2017; Chen et al. 2019) and of wind pollination (Culley et al., 2002) in temperate regions has been reported or suggested independently of mycorrhizal information. Likewise, AM-type trees are more frequently found in tropical regions and EcM-type trees in temperate regions (Gomes et al., 2019). The latitudinal gradients of seed dispersal and pollination may be the result of increased dependence on these interaction modes at the community and ecosystem scales through this differential dominance.

In conclusion, we found evidence of the joint evolution of mycorrhizal type, seed dispersal, and pollination mode in tree species. These findings strongly imply that mycorrhizal type shaped the evolution of seed dispersal and pollination modes, which have different yet physiologically and ecologically linked functions. The importance of such interactive effects highlights the need for more integrative studies to better understand how interactions shape the evolution of diversity in trees. In addition, our results highlight that most AM-associated trees, which are more common globally, are endozoochorous and biotically pollinated, and therefore may depend on mammals, birds, and insects for seed dispersal and pollination. The “empty forest syndrome” describes forests in which seed-dispersing animals are absent, their numbers decreased by hunting or human encroachment (Caughlin et al., 2015), and where aggregation of the same species is promoted (Wandrag et al., 2017). The aggregation of conspecific AM-type trees may increase negative PSFs in conspecific seedlings (Bennett et al., 2017; Tedersoo et al., 2020). Moreover, it has been reported that the quantity and species diversity of insects, including pollinators, is decreasing worldwide (Caughlin et al., 2015; Lister & Garcia, 2018; Sánchez-Bayo & Wyckhuys, 2019). Our findings emphasise the importance of protecting animals as seed dispersers and pollinators for the conservation of forest ecosystems. Most of the current forest conservation practices have focused on tree loss, such as deforestation (Alroy, 2017; Burivalova et al., 2019); these practices can be beneficial for preventing direct forest loss. However, our findings indicate that long-term conservation of forest ecosystems also requires the conservation of seed dispersers and pollinators in these ecosystems, as well as the conservation of a more comprehensive range of organisms that support their survival. Thus, conservation of forest ecosystems may require conservation activities focused not only on trees, but also on other organisms, with the goal of holistic long-term conservation.

## Author Contributions

Akira Yamawo (AY) and Misuzu Ohno (MO) developed the core idea. AY and MO performed the data analyses and wrote the paper. All authors contributed critically to the drafts and gave final approval for publication.

## ACKNOWLEDGEMENTS

We thank Yudai Okuyama and Hiroshi Ikeda for the helpful comments regarding the statistical analysis. AY thanks, Kiyoshi Ishida, Kenji Suetugu, Akihiro Kimura, and Tetsuro Yoshikawa for discussions during the initial stages of this project. This work was also supported by the Japan Society for the Promotion of Science (JSPS) Grant-in-Aid for Scientific Research (KAKENHI) (grant no. 18K19353 and 19H03295 to AY).

## Supplementary materials

**Supplementary Information** is available for this paper

Materials and Methods

Supplementary Text

Tables S1 to S5

---

**Table S1.** Results of the phylogenetically corrected Freeman–Tukey test for relationship between mycorrhizal type and seed dispersal mode.

| Mycorrhizal type | Seed dispersal mode | Observed phylogenetic species richness (PSR) | Expected phylogenetic species richness (PSR) | Freeman–Tukey deviate (FTD) | Freeman–Tukey significance test |
| --- | --- | --- | --- | --- | --- |
| AM | endozoochorous | 128.43526 | 93.06474 | 3.389978966 | $P < 0.01$ (+) |
| | anemochorous | 51.08221 | 82.349133 | -3.812830532 | $P < 0.01$ (–) |
|  | unspecialised | 22.61882 | 26.722419 | -0.771151045 | ns |
| AM+EcM | endozoochorous | 7.379251 | 10.38521 | -0.911162662 | ns |
|  | anemochorous | 9.247999 | 9.189444 | 0.097566252 | ns |
|  | unspecialised | 5.929395 | 2.981989 | 1.471862621 | ns* |
| EcM | endozoochorous | 6.061223 | 38.42578 | -7.318719596 | $P < 0.01$ (–) |
| | anemochorous | 65.209752 | 34.001384 | 4.507256651 | $P < 0.01$ (+) |
|  | unspecialised | 12.189693 | 11.033501 | 0.404951202 | ns |

AM: arbuscular mycorrhizal; EcM: ectomycorrhizal.
“+” Significantly higher and “–” significantly lower number of species than expected by the log–linear model with Bonferroni’s correction (FTD critical value = 2.305705); ns, not significant; \*no test because expected PSR < 5.

**Table S2.** Results of the phylogenetic Generalized Least Squares analysis of endozoochorous, anemochorous, and unspecialized seed dispersal distances. All data are log _10_-transformed before analysis.

|  | Maximum |  |  |  |  |  | Mean |  |  |  |  |  |
| --- | --- | --- | --- | --- | --- | --- | --- | --- | --- | --- | --- | --- |
|  | Endozoochorous |  |  | Unspecialized |  |  | Endozoochorous |  |  | Unspecialized |  |  |
| | $\lambda$ | *F*-value | *p*-value | $\lambda$ | *F*-value | *p*-value | $\lambda$ | *F*-value | *p*-value | $\lambda$ | *F*-value | *p*-value |
| Anemochorous | 0.58 | 5.44 | 0.02 | 0.73 | 16.28 | 0.0001 | 0.35 | 8.00 | 0.005 | 0.32 | 20.24 | <0.0001 |
| Unspecialized | 0.53 | 34.37 | <0.0001 | - | - | - | 0.42 | 27.46 | <0.0001 | - | - | - |

**Table S3.** Results of the phylogenetic Generalized Least Squares analysis of secondary seed dispersal distances and endozoochorous, anemochorous, and unspecialized seed dispersal distances. All data are log _10_-transformed before analysis.

|  | Maximum |  |  | Means |  |  |
| --- | --- | --- | --- | --- | --- | --- |
| | $\lambda$ | *F*-value | *p*-value | $\lambda$ | *F*-value | *p*-value |
| Endozoochorous | 0.315 | 62.12 | < 0.0001 | 0.565 | 94.19 | < 0.0001 |
| Anemochorous | 0.538 | 12.46 | 0.001 | 0.571 | 11.54 | 0.0018 |
| Unspecialised | 0.577 | 0.59 | 0.444 | 0.706 | 0.51 | 0.476 |

**Table S4.** Results of the phylogenetically corrected Freeman–Tukey test for relationship between mycorrhizal type and pollination mode.

| Mycorrhizal type | Pollination mode | Observed PSR | Expected PSR | FTD | Freeman–Tukey significance test |
| --- | --- | --- | --- | --- | --- |
| AM | Biotic | 145.24278 | 101.86939 | 3.933931571 | $P < 0.01$ (+) |
| | Wind | 45.98906 | 89.36245 | -5.296390431 | $P < 0.01$ (–) |
| AM + EcM | Biotic | 9.203612 | 12.10848 | -0.802871116 | ns |
|  | Wind | 13.526737 | 10.62187 | 0.894763163 | ns |
| EcM | Biotic | 7.765638 | 48.23416 | -8.17874215 | $P < 0.01$ (–) |
| | Wind | 82.780771 | 42.31225 | 5.203627492 | $P < 0.01$ (+) |

AM: arbuscular mycorrhizal; EcM: ectomycorrhizal.
“+” Significantly higher and “–” significantly lower number of species than expected by the log–linear model with Bonferroni’s correction (FTD critical value = 1.996799); ns, not significant.

**Table S5.** Results of the phylogenetically corrected Freeman–Tukey test for relationship between seed dispersal mode and pollination mode.

| Seed dispersal mode | Pollination mode | Observed PSR | Expected PSR | FTD | Freeman–Tukey significance test |
| --- | --- | --- | --- | --- | --- |
| Endozoochorous | Biotic | 107.44939 | 71.31273 | 3.860731941 | $P < 0.01$ (+) |
| | Wind | 24.68062 | 60.81728 | -5.593547003 | $P < 0.01$ (–) |
| Anemochorous | Biotic | 23.77434 | 65.55376 | -6.370626742 | $P < 0.01$ (–) |
| | Wind | 97.6853 | 55.90588 | 4.830189506 | $P < 0.01$ (+) |
| Unspecialised | Biotic | 24.76781 | 19.12505 | 1.249507397 | ns |
|  | Wind | 10.66756 | 16.31032 | -1.456970508 | ns |

“+” Significantly higher and “–” significantly lower number of species than expected by the log–linear model with Bonferroni’s correction (FTD critical value = 1.996799); ns, not significant.

